# Unpredictable hummingbirds: Flight path entropy is constrained by speed and wing loading

**DOI:** 10.1101/2020.08.11.246926

**Authors:** Ilias Berberi, Paolo S. Segre, Douglas L. Altshuler, Roslyn Dakin

## Abstract

Unpredictable movement can provide an advantage when animals avoid predators and other threats. Previous studies have examined how varying environments can elicit unpredictable movement, but the intrinsic causes of complex, unpredictable behavior are not yet known. We addressed this question by analyzing >200 hours of flight performed by hummingbirds, a group of aerial specialists noted for their extreme agility and escape performance. We used information theory to calculate unpredictability based on the positional entropy of short flight sequences during 30-min and 2-hour trials. We show that a bird’s entropy is repeatable, with stable differences among individuals that are negatively correlated with wing loading: birds with lower wing loading are less predictable. Unpredictability is also positively correlated with a bird’s overall acceleration and rotational performance, and yet we find that moment-to-moment changes in acceleration and rotational velocities do not directly influence entropy. This indicates that biomechanical performance must share an underlying basis with a bird’s ability to combine maneuvers into unpredictable sequences. Contrary to expectations, hummingbirds achieve their highest entropy at relatively slow speeds, pointing to a fundamental trade-off whereby individuals must choose to be either fast or unpredictable.

## INTRODUCTION

One of the most effective ways to avoid predators and other threats is to escape by moving in an unpredictable way [1–3]. Previous studies have tested how environmental factors, such as the presence of a predator, can cause individuals to temporarily become less predictable [4–9]. Experiments have also shown that unpredictability can be advantageous, both to escaping animals [10] and those in pursuit of prey [11]. However, it is not yet known whether some individuals are inherently more unpredictable than others, as a result of inter-individual differences in the underlying control systems. These inter-individual differences are necessary for selection to act on unpredictable behavior [3,12].

Biomechanical traits, such as morphology and muscle composition, are one potential driver of individual differences in unpredictability [7,13]. Morphology strongly predicts the locomotor capabilities of individuals [14,15] and covaries with evolved behavioral differences across species [9,16–18]. An animal’s morphology and muscle composition also determine limb kinematics that influence whole-body maneuvers [19–21]. These maneuvers are combined as sub-units to generate longer sequences of behavior that may vary in higher-order properties such as unpredictability [9] (Fig. 1). The available data suggest two non-exclusive hypotheses for the relationship between morphology, biomechanical traits, and unpredictable behavior. The first hypothesis, which we refer to as the facilitation hypothesis, is that enhanced power-generation may be associated with greater unpredictability [9,22,23]. This can occur if high-performance maneuvers directly confer unpredictability, or if there is a shared underlying basis for maneuverability and unpredictability [17]. For example, more maneuverable individuals may have sensory or neuromuscular advantages that also allow them to combine behavioral sub-units in more unpredictable ways. The second hypothesis, which we refer to as the trade-off hypothesis, is that power generation may be negatively associated with unpredictability [7]. This can occur if high-performance maneuvers restrict an animal’s ability to vary its subsequent trajectory [24], due to constraints of inertia and/or sensory and neuromuscular processing. Testing these broad hypotheses is challenging, because it requires tracking individuals that are free to perform diverse maneuvers and combine them in different ways.

**Figure 1.**
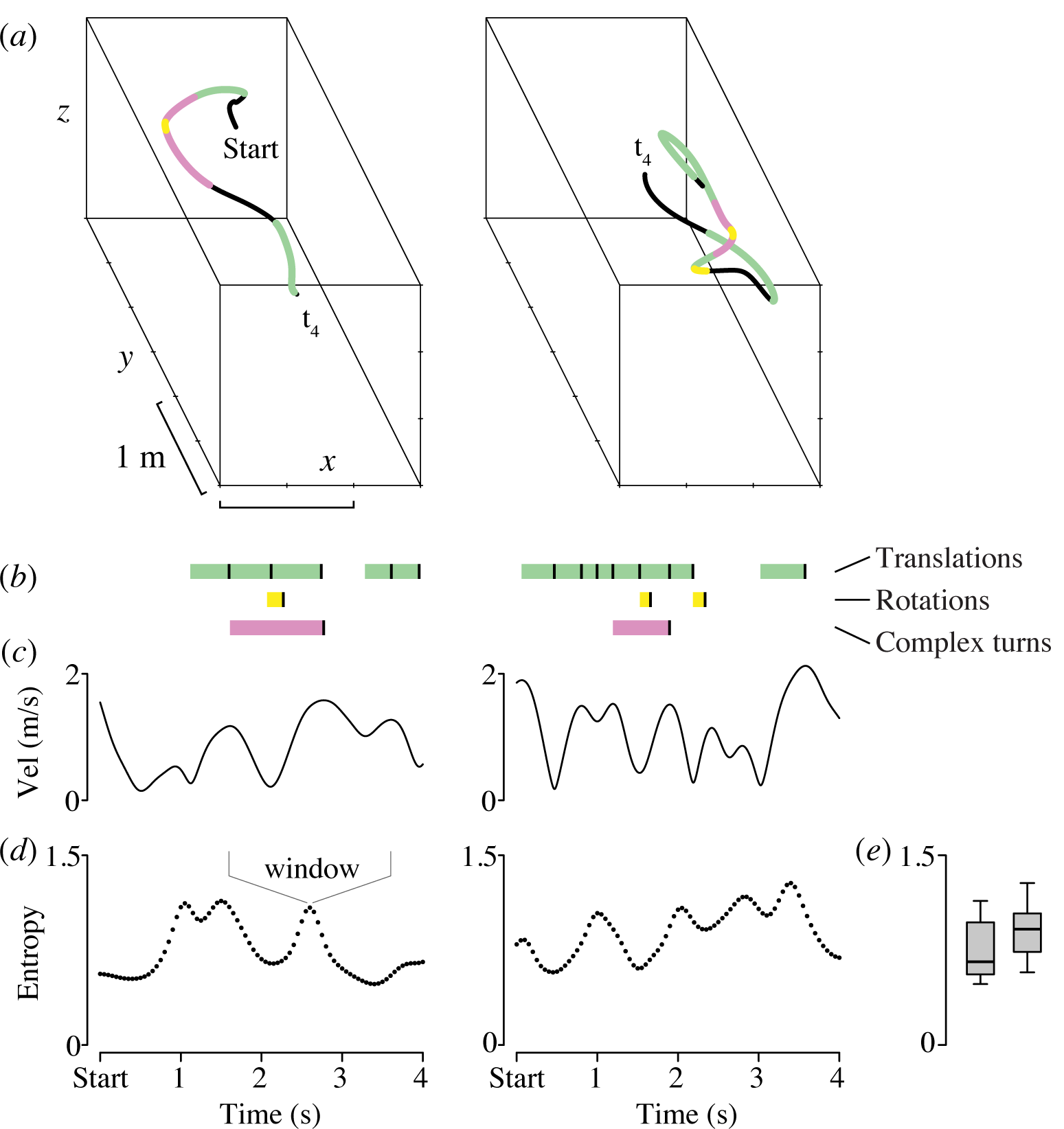
Hummingbirds combine simple maneuvers into longer flight sequences. (a) Examples of two 4-s sequences of hummingbird flight, sampled at 200 fps. The highlighted points ‘Start’ and ‘t4’ correspond to the x-axis tick marks on the time series below in (b-d). (b) The timing of translations, rotations, and complex turn maneuvers. The black vertical lines denote the ends of each maneuver. (c) Total velocity vs. time. (d) Positional entropy vs. time, defined as the difficulty of predicting a bird’s location during forward flight. Each entropy value is calculated from the surrounding 2 s window. (e) Entropy boxplots for the two illustrated sequences. The sequence on the left is more predictable than the one on the right (average entropy 0.74 vs. 0.90).

Hummingbirds provide an opportunity for studying unpredictability because they are extremely agile in three dimensions, excellent at evading threats, and their survival and mating success often depend on locomotor performance [25]. One of the primary reasons that hummingbirds are so agile is their extremely high wingbeat frequencies (up to 80 Hz), which generate rapid forces on both the upstroke and downstroke [26,27]. This allows hummingbirds to perform a wide range of aerial maneuvers that can be combined into complex sequences of behavior during pursuit and escape [28–30] (Fig. 1b, Table 1). Previous studies of flying hummingbirds have shown that the kinematics of the wings and body are influenced by inter-individual variation in body mass, flight muscle mass, and wing loading [31–33]. More recently, these morphological traits have also been mapped to individual differences in the use and performance of different maneuvers defined in Table 1 [28,29,34]. For example, flight muscle capacity is correlated with enhanced performance on multiple translational, rotational, and turning performance metrics. Reduced wing loading is also associated with enhanced performance of aerial maneuvers. Finally, body mass variation within species has been shown to have a detrimental effect on performance. These previous studies provide a foundation for understanding agility in avian flight, making hummingbirds a tractable system for investigating the mechanistic basis of unpredictable movement.

**Table 1.**
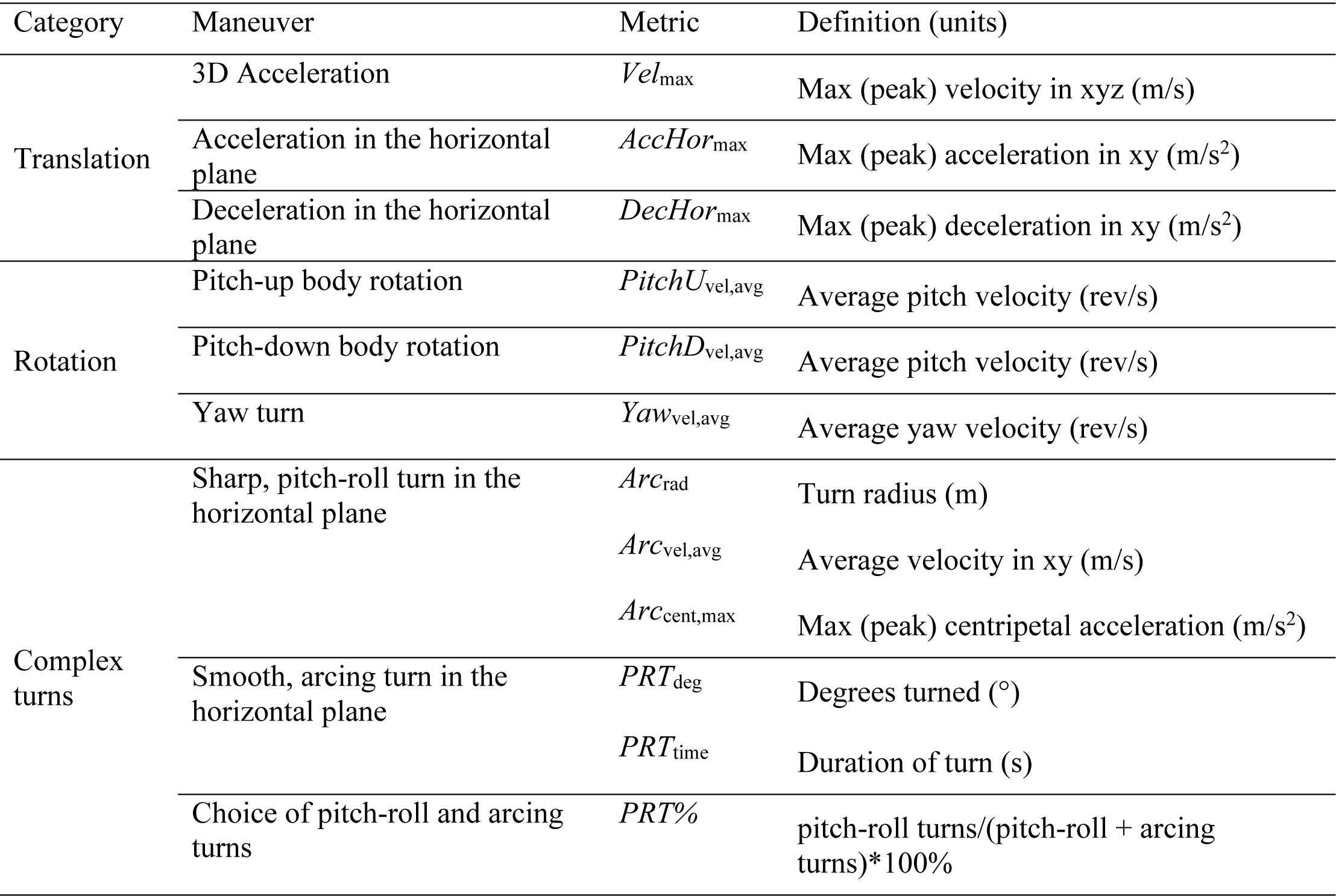
Hummingbird flight maneuvers and performance metrics. For detailed descriptions on all maneuvers see [28].

Here, we test whether variation in unpredictability is related to biomechanical traits and maneuvering performance. We use data from previous studies of hummingbirds where the individual birds were tracked extensively in a large chamber and were free to choose their sequence of flight behaviors. The birds often flew while motivated by the variable presence of the human experimenter or other startling stimuli. This method allowed us to record a range of rapid and ballistic maneuvers, such as those shown in Fig. 1. To evaluate whether unpredictability may be driven by intrinsic among-individual differences, we used positional entropy to quantify unpredictability [8–10] and tested whether it was repeatable among-individuals. Next, we used a comparative study of 207 individuals from 25 species to evaluate whether unpredictability differs among species and whether it is related to inter-individual differences in morphology and maneuvering performance. According to the facilitation hypothesis, we expected more unpredictable individuals to use higher-performance translations, rotations, and complex turns, whereas we had the opposite expectation under the trade-off hypothesis. Under the facilitation hypothesis, we also expected a bird’s burst muscle capacity to be positively associated with unpredictable behavior, and its body mass and wing loading to be negatively associated with unpredictability. Additionally, we examined wing aspect ratio, a trait that is associated with energetic efficiency and with the use of certain turns in hummingbirds [29,35]. If unpredictability requires energetic efficiency, then we expected individuals with higher aspect ratio wings to be more unpredictable. Next, to evaluate whether the use and performance of specific maneuvers directly influences unpredictability, we used a within-individual analysis. This analysis tested whether moment-to-moment changes in a bird’s predictability were associated with specific maneuvers. To provide context for these results, we also used a simulation to test how the speed of behavior theoretically influences unpredictability.

## MATERIAL AND METHODS

### (a) Flight trials and tracking system

We used tracking data on wild-caught hummingbirds that were recorded in two previous studies: (1) a repeatability study in which the same 20 individuals were assayed in multiple flight trials [28], and (2) a comparative study that assayed 207 individuals from 25 species [29]. Each study captured individuals at multiple sites (3 sites for repeatability and 4 for comparative, described in [28,29]). To assay flight behavior, the birds were tracked during flight trials in a large 3 m x 1.5 m x 1.5 m chamber where they were motivated by a variety of environmental stimuli, including the threat of the human experimenter. In 32 of the trials in the repeatability study, a competing bird was also present. Flight trials lasted for 2 hours in the repeatability study, and 30 min in the comparative study. Flight in the chamber was recorded with four or five synchronized cameras (GE680, Allied Vision Technologies, Stadtroda, Germany) filming at 200 fps. An automated tracking system was used to determine the body position and orientation of the birds in each frame [36]. The output of the tracking system was smoothed using a forward-reverse Kalman filter and ground-truthed by re-projecting the smoothed data onto the videos, to verify that it matched the movements of the animals [28,34,36].

### (b) Quantifying unpredictability

Some of the earliest work conceptualizing unpredictability focused on phenomenological descriptions [3]. Modern computational approaches provide a standardized way to quantify unpredictability using information theory, allowing individuals and species to be compared [8– 10]. We used this approach to quantify the unpredictability of hummingbird flight paths based on positional entropy, defined as the ability to predict an animal’s future position given its past trajectory [37]. Large values of positional entropy correspond to highly unpredictable movement and are indicative of multiple dynamic changes in speed and heading. Positional entropy is calculated based on a series of spatial coordinates in one, two, or three dimensions (see [37] for equations). Mathematically, the computation ignores the scale of the spatial units, such that a sequence of behavior with a given spatial configuration (like a perfect circle with constant velocity) would have the same entropy, regardless of whether it has a 2 cm or 2 m radius. However, positional entropy is highly influenced by the temporal duration of the sequence: as the duration gets smaller, the positional entropy asymptotically approaches 0. Therefore, when using this metric for comparison, it is important to fix the temporal scale to a single duration that can be applied to all individuals.

Because positional entropy is a sequence-based metric, it is challenging to define the start and end points. In our hummingbird free-flight trials, the natural transitions between flight modes and maneuvers were highly variable. We therefore quantified positional entropy using a sliding window approach [38]. We chose a 2-s duration for each window (Fig. 1d), because this duration was long enough to contain multiple maneuvers defined in Table 1 (average 4 ± 2 SD), while also occurring many times within each trial. We identified each window at 50 ms intervals after the start of the trial. Because these sequences are overlapping and consecutive, they are not independent. All three-dimensional positional entropy calculations and further statistical analyses were performed in R (version 3.6.2) [39].

An important consideration when calculating entropy is that even when the animal is stopped, subtle movements of its body parts or unsmoothed tracking error can produce very high entropy values, because the calculation is normalized on a relative spatial scale. In any study of whole-body movement, it is therefore important to limit the analysis of positional entropy to sequences when the spatial changes are driven by whole body movement (as opposed to body parts). We used a threshold criterion whereby entropy was only calculated for sequences when the hummingbird’s total velocity was maintained above 0.1 m/s throughout all frames, to limit our analysis to forward flight.

### (c) Repeatability and species differences

To determine if unpredictability differs among individuals, we used data from 20 Anna’s hummingbirds (*Calypte anna*) in the repeatability study. All individuals were assayed once on their own, and 1-2 times in the presence of a competitor, as part of a previous experiment to test whether competitor presence affected maneuvering performance [28]. For the present study, we calculated a single trial-average entropy for each bird in a given trial (n = 52 trials). We used a mixed-effects model in the “lme4” package [40] with bird ID as the random effect to compute the repeatability of trial-average entropy [41]. The model controlled for site (3 levels) and competitor presence (present vs. absent) as the fixed effects. To determine if species differ in unpredictability, we also computed repeatability (at the species level) of trial-average entropy in the comparative study, controlling for site (4 levels; n = 207 individuals from 25 species) [28,29]. We determined the 95% confidence intervals of repeatability estimates by parametric bootstrapping with 10,000 iterations using the “bootMer” function in lme4.

### (d) Maneuvers, morphology, and entropy

To determine how a bird’s maneuverability was related to its unpredictability, we analyzed trial-average entropy in relation to 12 maneuvering performance metrics that were previously shown to be repeatable [28] and correlated with each other [29]. A brief description of each metric is provided in Table 1 [28,29]. Although the maneuvers in Table 1 are not a complete list of what hummingbirds can perform, they represent a simple set of behavioral units that are shared by all individuals studied. We first evaluated the association between an individual’s trial-average entropy and each performance metric by fitting a separate phylogenetic regression model for each metric, with entropy as the response variable and the performance metric as the predictor. Models were run in the Bayesian “MCMCglmm” package [42] using the most recent multilocus phylogeny from [43], and we included field site as a fixed effect to account for potential site differences.

Next, we evaluated which morphological and muscle traits best explain an individual’s overall unpredictability. Measures of body mass, wing loading, and aspect ratio were taken from the same individuals in the maneuvering trials [29]. We obtained estimates of species-average muscle capacity based on previous studies that used a transient load-lifting assay [29,32,44]. This analysis was also run as a Bayesian phylogenetic regression in MCMCglmm, with the following predictors: individual body mass, wing-loading, and aspect ratio, species muscle capacity, and field site.

All MCMCglmm models were run with uninformative priors for Gaussian data. Each chain was run for 500,000 iterations after a burn-in period of 3,000 and the posterior results were thinned to include every 500^th^ sample. We verified that autocorrelations in the posterior samples were < 0.1 and confirmed that models had converged by checking the Gelman-Rubin statistic for two independently seeded chains. We report standardized effect sizes as the posterior mean ± 95% credible intervals.

### (e) Within-individual analysis and simulated effect of speed

Using data from the comparative study, we tested whether particular maneuvers could explain moment-to-moment changes in entropy. We computed a series of additional metrics for each 2 s sliding window: (i) travel velocity, as the sum of the Euclidean distances travelled between each frame, divided by the time (2 s), (ii-v) the total number of horizontal translational maneuvers, rotational maneuvers, pitch-roll turns, and arcing turns performed in the time window, and (vi-vii) the average performance on any translational maneuvers (m/s^2^) and any rotational maneuvers (rev/s) performed in that window. For (vi), *AccHor*_max_ and *DecHor*_max_ were combined, and *PitchU*_vel,avg_, *PitchD*_vel,avg_, and *Yaw*_vel,avg_ were combined for (vii), because most 2 s windows did not include a maneuver of each type (Table S1).

We fit a linear mixed-effects model in the “nlme” package [45] to analyze entropy for each 2 s window as a function of the predictors (i-vii) above. The model included nested random effects of species, individual ID, and flight bout to account for non-independence at these three levels. A flight bout included any sequential 2-s windows; after a gap > 50 ms, a new bout was assigned. To account for strong temporal autocorrelation, the model included the corAR1 autocorrelation structure [46]. The sample size for this analysis was limited to windows that had at least one translation and one rotation (n = 251,545 windows for 205 individuals). We also checked models that dropped average translational and/or rotational performance to verify that the conclusions for the other predictors were unchanged when analyzing the full dataset.

We used a simulation to test the theoretical effect of the speed on entropy. The simulation assumed that individuals in the 20 solo trials of the repeatability study could perform the same movement geometry either twice as fast, or twice as slow. To simulate an increase in speed (2x faster), we removed every other recorded frame throughout each trial. This method retained the same spatial geometry of the flight paths, but performed it in half the time (see Fig. S1 for an example). To simulate a decrease in speed (2x slower), we added a new frame between each original frame and interpolated the bird’s coordinate values. This treatment also retained the same spatial geometry, but performed it in twice as much time (Fig. S1). We recalculated trial-average entropy based on sliding 2 s windows in the newly simulated timeframes. To analyze the simulation outcome, we fit a linear mixed effects model in the “lme4” package, with trial-average entropy as the response variable, speed condition (original, slower, or faster) as the fixed effect, and individual identity as a random effect.

## RESULTS

### Repeatability and species differences

Some hummingbirds are consistently more unpredictable (higher entropy) than others, as indicated by moderate and significant repeatability (Fig. 2a; R = 51%, 95% CI = 20–77%, p = 0.002). We did not detect a significant effect of competitor presence on entropy (Table S2). Our comparative analysis also indicated that species differ in unpredictability, with inter-species differences accounting for 24% of the total variance in entropy (Fig. 2b, 95% CI = 6–42%, p = 0.002).

**Figure 2.**
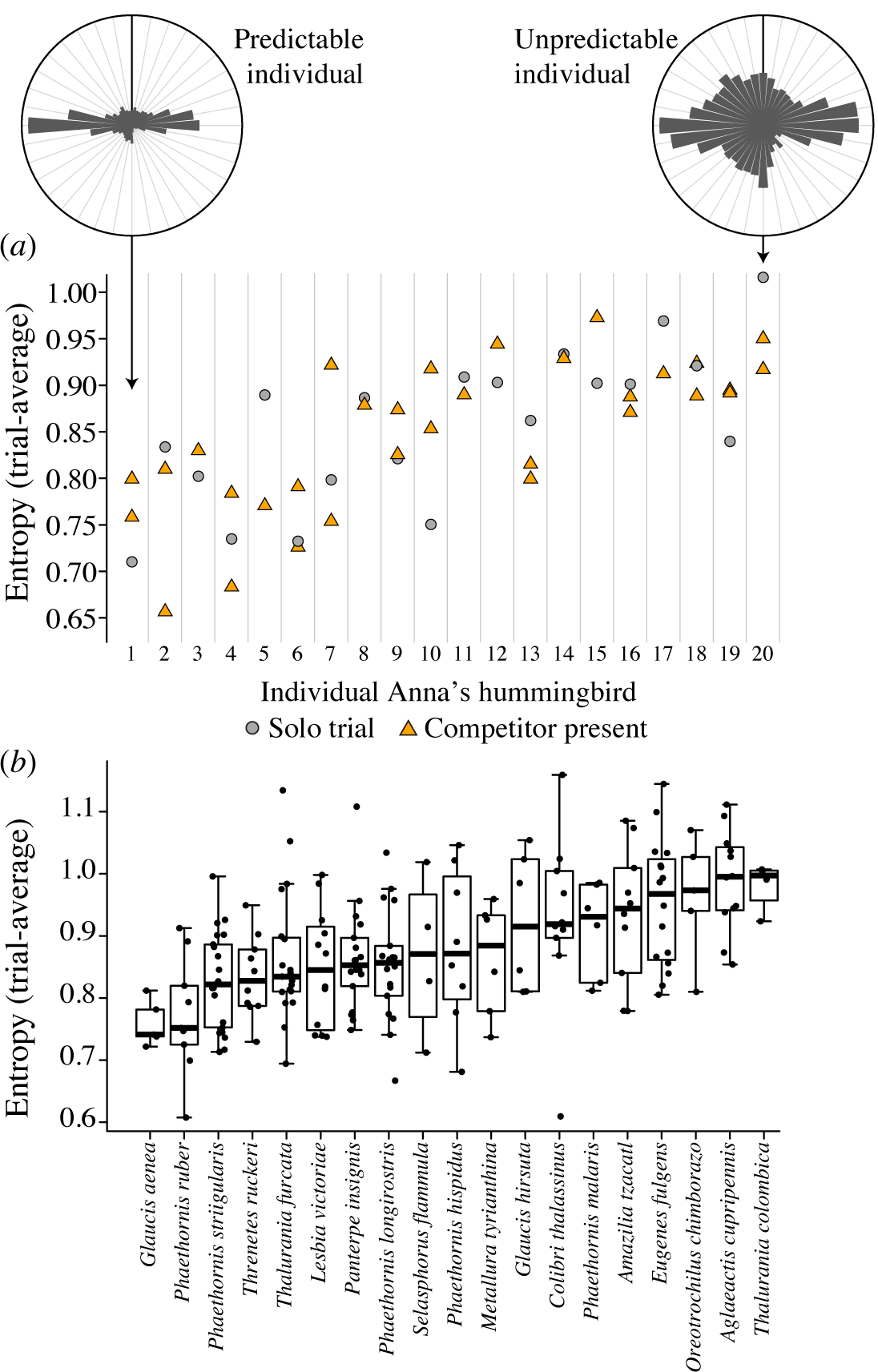
Individuals differ in unpredictability. (a) Trial-average entropy was repeatable for 20 male Anna’s hummingbirds tested in 2-3 trials (each bird = one column, sorted by individual mean). Circular histograms show headings for the most predictable individual (left) and the most unpredictable individual (right), from a sample of sequences at their most common entropies (vertical line = 0° heading). Headings were more variable for the high-entropy individual. (b) Species differ in unpredictability. Only the 19 species that had >2 individuals are shown.

### Maneuvers, morphology, and entropy

The most unpredictable birds used lower average peak velocities (*Vel*_max_), but higher average peak accelerations and decelerations (*AccHor*_max_, *DecHor*_max_; Fig. 3a, Table S3). These contrasting effects were surprising, because individual variation in *Vel*_max_ is positively correlated with *AccHor*_max_ and *DecHor*_max_ performance [29] (Fig. S2). The most unpredictable hummingbirds also used faster rotational maneuvers (pitch-up, pitch-down and yaw; Fig. 3a, Fig. S3). Most of the features of the arcing and pitch-roll turns were not strongly related to entropy (Table S3). The exceptions were *PRT*_time_ and *PRT*%: the most unpredictable hummingbirds tended to perform slightly slower pitch-roll turns and they also used pitch-roll turns in their flight trials more often (Fig. 3a).

**Figure 3.**
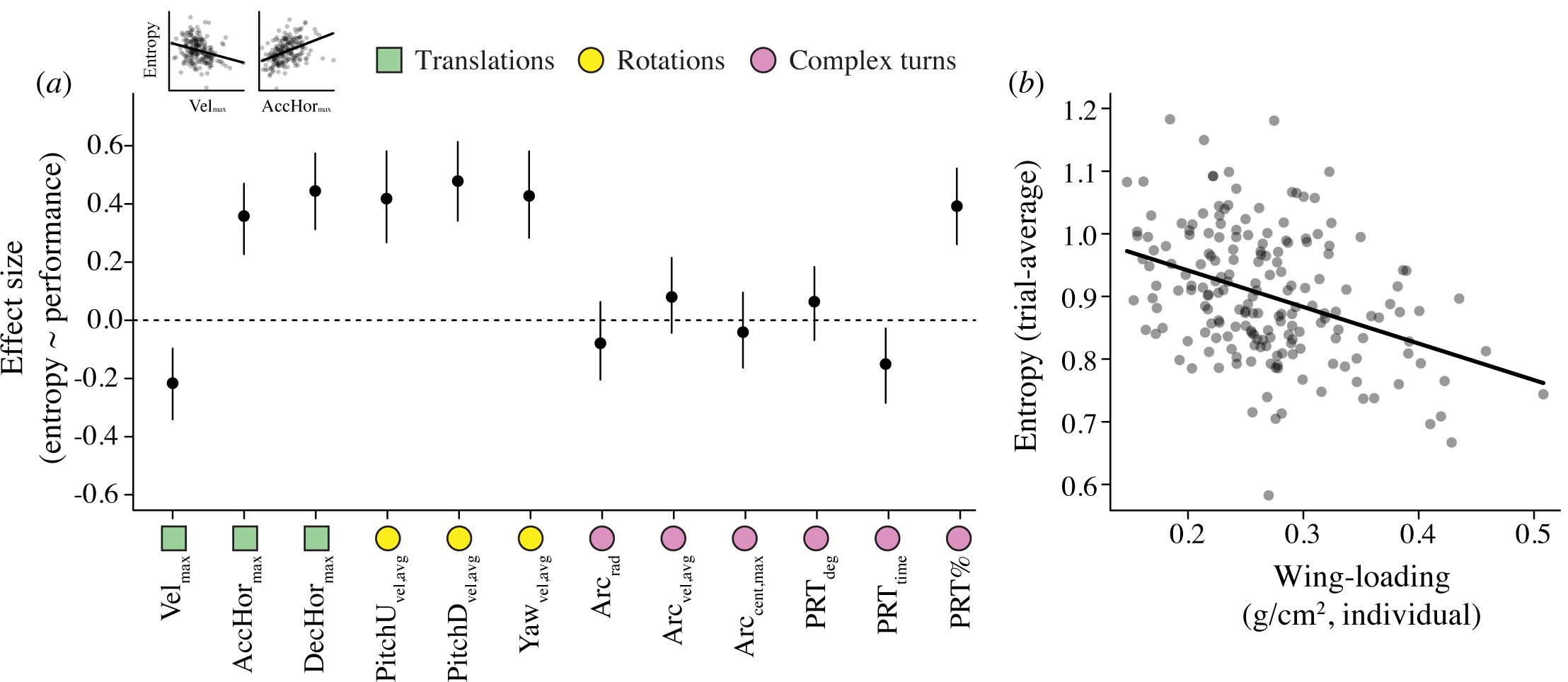
Unpredictable behavior is associated with an individual’s maneuvering performance and wing morphology. (a) Slope estimates ± 95% CI for the relationship between a bird’s average entropy and maneuvering performance metrics. See Table 1 for definitions. Positive slopes indicate that higher performance is associated with greater unpredictability (n = 207 individuals from 25 species). All predictors and the response variable were centered and standardized, so that the values plotted here are comparable effect sizes. Inset examples show that entropy is negatively related to Vel_max_ but positively related to AccHor_max_. (b) Unpredictable individuals have lower wing loading (i.e., more wing area per unit body mass). The y-axis shows partial residuals for entropy after accounting for other predictors in the analysis. See Tables S3-S4 for details.

Examining morphology, the strongest predictor of entropy was wing-loading, such that the most unpredictable hummingbirds had lower wing-loading than others (Fig. 3b, Table S4). Although there was a weak positive relationship between entropy and species muscle capacity, the posterior credible interval included 0. Body mass and aspect ratio did not explain variation in entropy.

### Effect of speed and the within-individual analysis

All else being equal, if an animal is able to perform the same sequence of behavior twice as fast, its entropy is expected to increase, whereas entropy is expected to decrease for an animal moving more slowly (Fig. 4a; Table S5). Examining variation in the actual hummingbird trajectories, the best predictor of moment-to-moment changes in entropy was travel velocity. However, contrary to the simulation result described above, hummingbirds were more unpredictable when moving slower (Fig. 4b, Table S6). Entropy within a given flight sequence was also positively associated with translational performance, but not rotational performance (Fig. S4). The lack of rotational effect was surprising, because both translational and rotational performance were strongly associated with inter-individual differences in entropy (Fig. 3a and S4). Furthermore, the association between entropy and translational performance was much stronger at the among-individual level than it was within individuals (3-to-4 fold difference in effect size; Fig. S5). High entropy sequences also had more rotations and complex turns, but fewer translations (Table S6). However, while these effects were statistically significant, the performance and use of specific maneuvers could only explain 0.4–1.7% of the variation in unpredictability within-individuals (Table S6). In contrast, the constraint of travel velocity explained much more of the variation in entropy (R^2^ near 9%).

**Figure 4.**
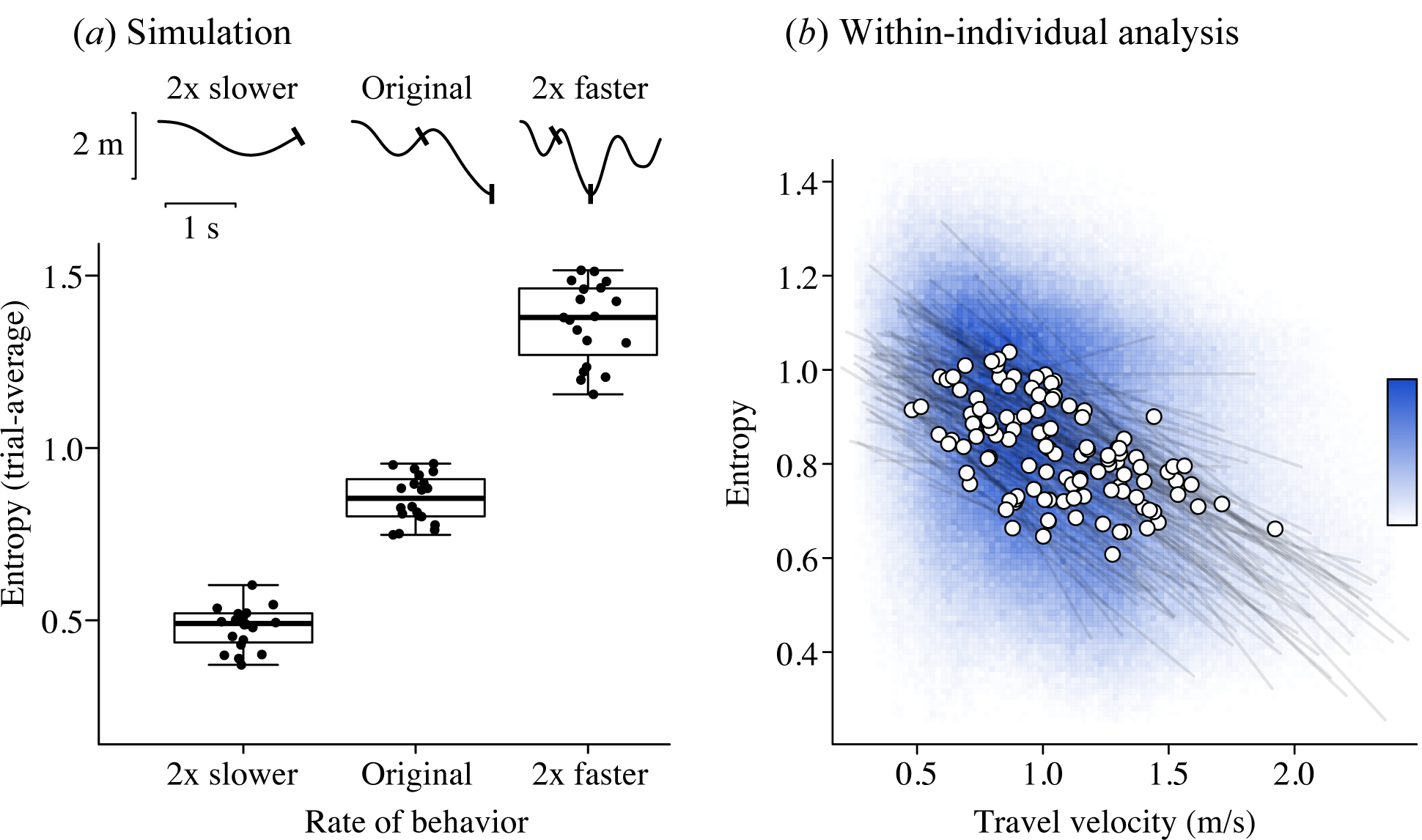
The trade-off between speed and unpredictability. (a) A simulation tested the effect of performing the same behavioral sequence 2x slower or 2x faster. The traces above (a) show position (y-axis) vs. time (x-axis), with tick-marks denoting time points from the original, to show how time is either compressed or expanded (Fig. S1). The boxplot in (a) shows that in principle, faster flight is expected to be less predictable. (b) However, in reality, hummingbirds are more predictable at higher speeds, owing to performance and/or sensory constraints. The blue color is used to represent the frequency of entropy data points per pixel (n = 732,855 values from 207 birds; the darkest blue indicates 281 values per pixel; white pixels have 0 values). Grey lines show model predictions for each individual’s entropy as a function of speed and controlling for temporal autocorrelation. White dots show the predicted values at a bird’s median speed. The average number of entropy values per bird was 3,540. For clarity, this plot only shows individuals with at least 2,000 entropy values. See Table S5.

## DISCUSSION

Our studies of hummingbirds reveal that some individuals and some species are consistently more unpredictable than others (Fig. 2). This raises possibility that unpredictability may be heritable [12,47] and correlated with other intrinsic traits. In our analysis of trial-average entropy, we found strong associations between an individual’s unpredictability and biomechanical performance: the most unpredictable hummingbirds used higher-performance accelerations, decelerations, and rotational velocities; they also tended to use pitch-roll turns more often (Fig. 3a). Therefore, hummingbirds with greater capacity for power generation during their maneuvers also move in more unpredictable ways, consistent with the facilitation hypothesis. Given these findings, it was not surprising that a hummingbird’s wing-loading was also related to flight path entropy: larger-winged hummingbirds are less predictable (Fig. 3b). Previous research has shown that species with lower wing-loading also perform faster rotations and more sharp turns [29], both of which were also associated with an individual’s overall unpredictability in our analysis. However, contrary to the facilitation hypothesis, we found that unpredictability was not related to burst muscle capacity, one of the most important traits that determines power generation for flying hummingbirds [28,32,44].

Given the strong associations between an individual’s overall maneuverability and entropy (Fig. 3), we expected to find that within individuals, hummingbirds would be less predictable when using fast translational and rotational maneuvers. Surprisingly, we found that moment-to-moment changes in rotational performance were virtually unrelated to entropy, and moment-to-moment changes in translational performance explained less than 2% of the variation in entropy within individuals (Fig. S4 and Table S6). This indicates that high-performance maneuvers make little to no direct contribution to unpredictability on the scale of our assay. Taken together, these findings imply that among-individual differences in unpredictable flight and high-performance maneuvers must share an underlying basis in some other trait(s) [17]. Possible candidates include a bird’s sensory capacities, neuromuscular performance, processing speed, body condition, energy budget, risk tolerance, or skill and expertise [48–50]. An important next step is to determine why we see strong inter-individual correlations between performance and unpredictability, and how these behavioral phenotypes and their underlying systems co-evolve [17,51].

The best predictor of moment-to-moment changes in unpredictability was a bird’s speed. Contrary to theoretical expectations, hummingbirds were more unpredictable when flying slowly (Fig. 4). This tradeoff was also observed among individuals: the most unpredictable hummingbirds used high accelerations while maintaining relatively low velocities (Fig. S2), pointing to a fundamental tradeoff between speed and entropy. We propose that flying fast restricts an animal’s ability to vary its subsequent trajectory, due to constraints of inertia and/or sensory and neuromuscular processing. These results are similar to ones found in fish [7] and mammals [24,52], suggesting that this trade-off may be a fundamental property of behavioral control systems.

Overall, we found support for both facilitation and trade-off hypotheses, indicating that the relationship between biomechanics and unpredictability is nuanced. In hummingbirds, speed trades-off with unpredictability, and yet individuals differ in traits (such as relative wing size) that allow some individuals to operate at a higher level of maneuvering performance and unpredictability. Given that the maneuvers themselves do not directly contribute to entropy variation in hummingbirds, our results raise new questions about the mechanisms underlying among-individual differences [17,49,53]. Based on our results, we propose that maneuvering performance is associated with a broader biomechanical-behavioral phenotype that also determines unpredictability [9]. Testing the basis of the proposed biomechanical-behavioral phenotype will require high-throughput approaches that capture variation across multiple scales, from sub-units of locomotion to whole sequences of behavior [38]. In addition to the biomechanical and behavioral perspectives, unpredictability carries important fitness consequences [10,24]. Future research should explore the intrinsic causes of unpredictability and its tradeoffs using predator-prey and pursuit contexts, where each interaction carries a measurable fitness component [13].

## ETHICS STATEMENT

All research was approved by the Institutional Animal Care and Use Committee of the University of California, Riverside, the University of British Columbia Animal Care Committee, the Ministerio de Agrigultura of Peru, the Ministerio de Ambiente y Energia of Costa Rica, and the Ministerio del Ambiente of the Pichincha province of Ecuador.

## DATA ACCESSIBILITY

All data and R scripts are available at: https://figshare.com/s/c1c07cd0dc0b8e9a67b1 The repository will be made public when the final version of the study is published.

## AUTHOR CONTRIBUTIONS

All authors designed the study. PSS collected the data. RD and IB analyzed the data and wrote the manuscript. All authors edited the manuscript.

## ACKNOWLEDGEMENTS

We thank the Los Amigos and La Selva Biological Stations, La Georgina restaurant, Hacienda Guaytara, Andrew D. Straw, and numerous field technicians and colleagues who contributed to references [28,29,34].

