## Supplementary Information for "Unpredictable hummingbirds: Flight path entropy is constrained by speed and wing loading"

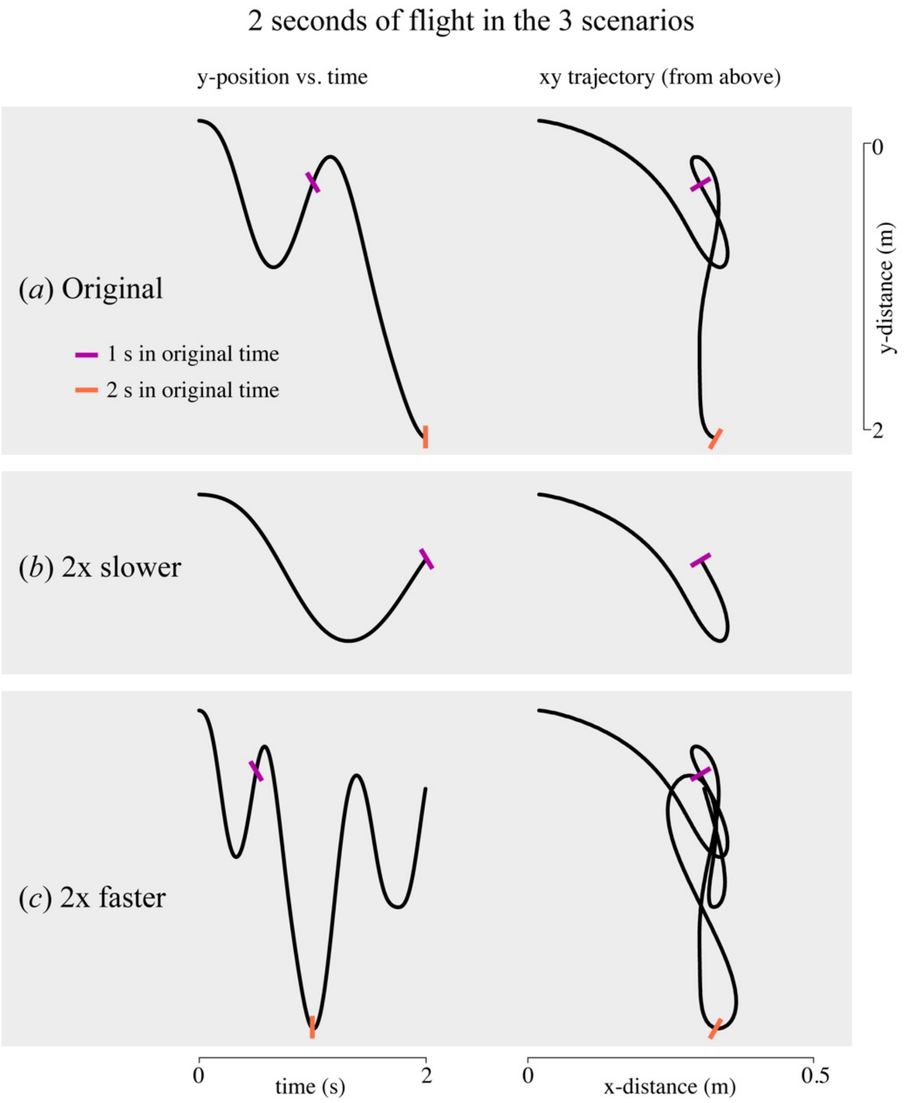

**Figure S1. Simulated change in the speed of hummingbird behavior.** Each example shows 2 s segment of hummingbird flight. In (a), we see 2 s of observed flight ('Original'). The left column (a) shows position in the y-axis vs. time. The right column (b) shows a view of the same trajectory from above, looking at the xy-plane. To simulate a decrease in speed (b; '2x slower'), time is expanded so that fewer displacements are performed per unit time. To simulate an increase in speed (c; '2x faster'), time is compressed, so that more displacements are performed per unit time. The colored markers show correspondence with the original timeframe. Note that for visibility, the scales of the x-distance and y-distance axes are distinct, and the scale for y-distance applies to all panels of the figure.

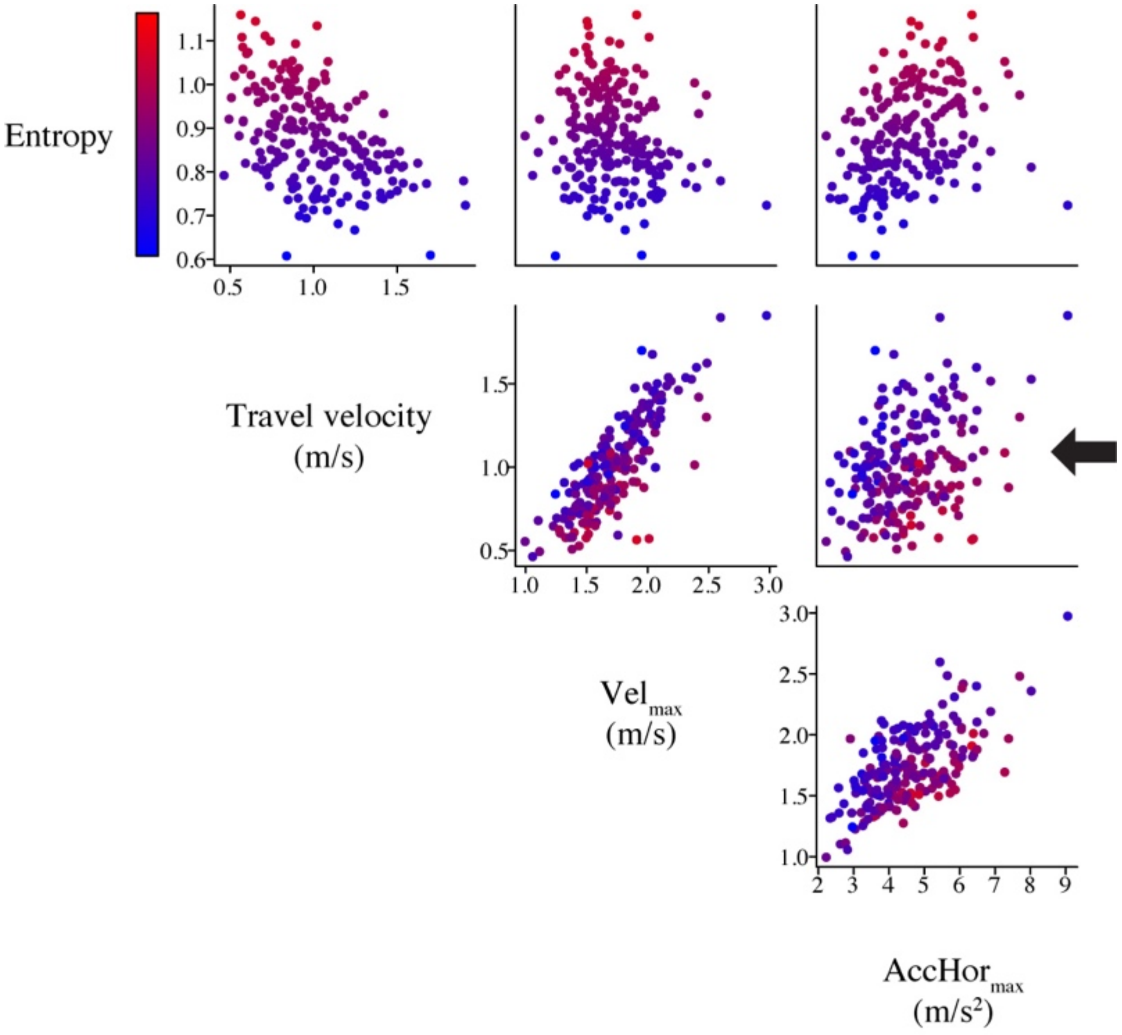

**Figure S2. Scatterplot matrix showing the relationship between a bird's trial-average unpredictability, speed, and acceleration performance.** Each point represents the trial-average for a single bird (n = 207 birds from 25 species). Points are colored by an individual's entropy, according to the scale shown at the top left (red-colored datapoints = high entropy individuals). A bird's trial-average travel velocity is computed from the total distance travelled per sliding window divided by time. Average $Vel_{max}$ is calculated from the peak total velocities of horizontal acceleration maneuvers. Average $AccHor_{max}$ is calculated from the peak horizontal accelerations during those maneuvers. Even though speed and acceleration are positively correlated, the most unpredictable individuals used relatively high accelerations, but maintained relatively low velocities, as shown in the plot of $Vel_{max}$ vs. $AccHor_{max}$ on the middle right.

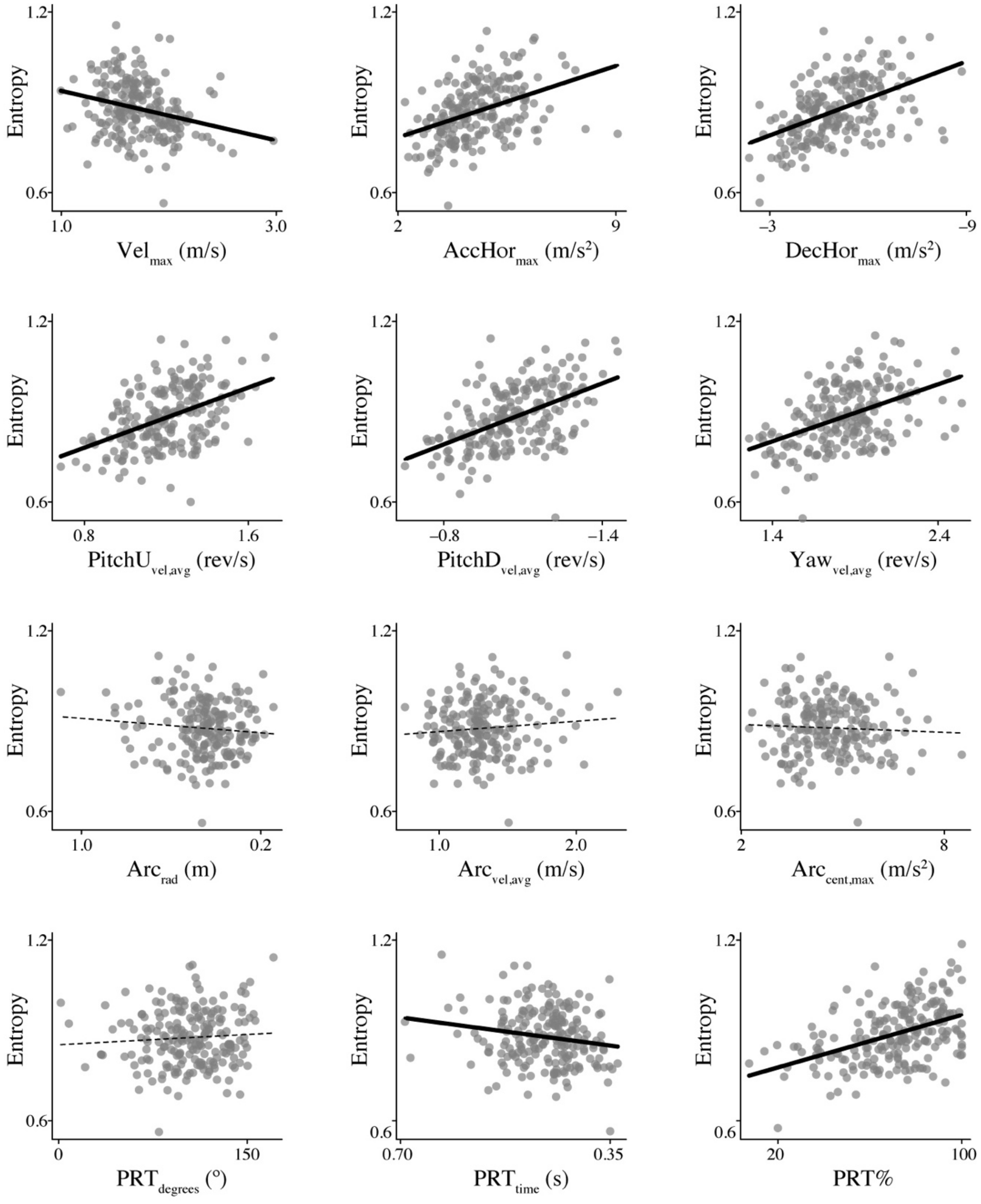

**Figure S3. Relationships between a bird's unpredictability and maneuvering performance.** Each point represents a single bird (n = 207 birds from 25 species). The y-axis shows the partial residuals for trial-average entropy, after accounting for other factors. Solid fit lines are used if the credible intervals of the posterior slope estimate in the phylogenetic regression excluded 0. Dotted lines indicate that the posterior distribution of the slope estimate broadly overlapped 0 (Table S3). See Table 1 in the main text for definitions of each performance metric.

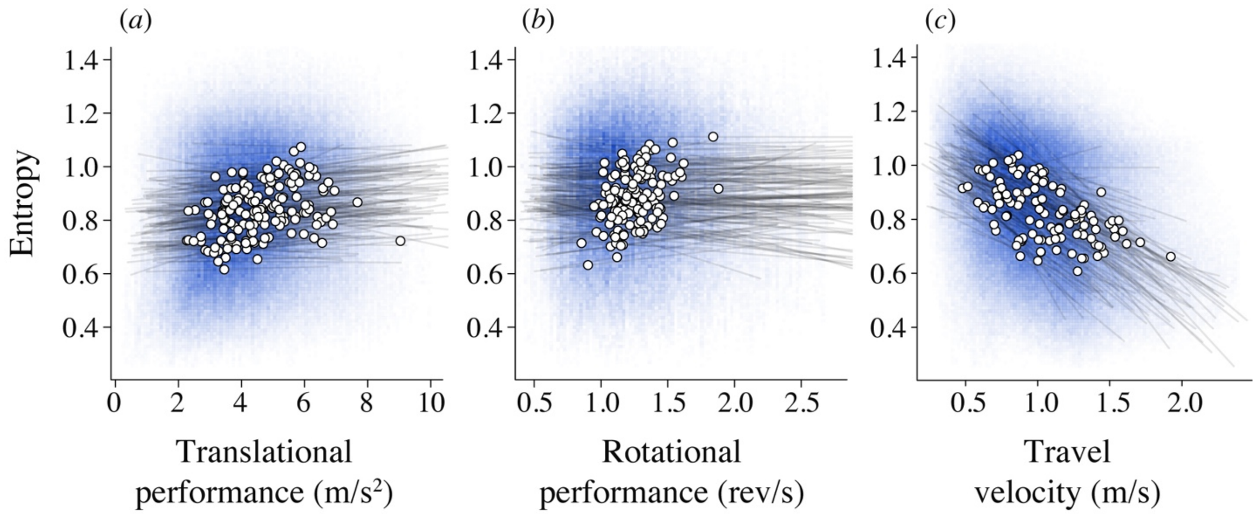

**Figure S4. A within-individual analysis tests the direct effects of maneuvering performance on unpredictability.** Each plot (a-c) shows one of the predictors from Table S6; note that (c) is reproduced from Fig. 4 in the main text for comparison. The blue color is used to represent the density of entropy data points per pixel (n = 732,855 values from 207 birds). The white dots and grey lines are used to visualize the among- and within-individual effects, respectively. Each white dot represents an individual's overall average and the gray lines show the model predictions within individuals. Although a bird's average entropy is positively correlated with its overall translational and rotational performance, the relatively flat slopes of the within-individual lines in (a-b) demonstrate that performance of these maneuvers does not directly contribute to unpredictability. In contrast, travel velocity (c) is negatively related to unpredictability both among and within individuals. For clarity, only individuals with at least 500 entropy vs. performance values are plotted in (a-b).

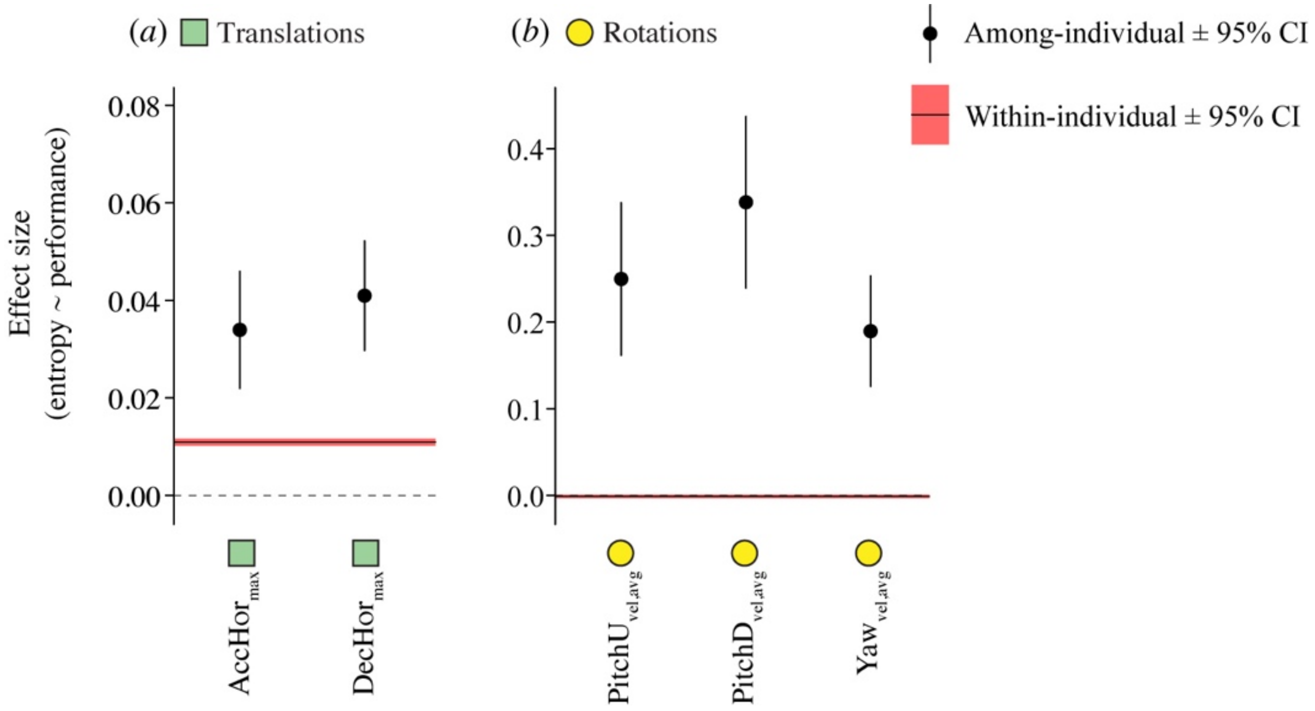

**Figure S5. Comparison of among- and within-individual effect sizes for entropy as a function of maneuvering performance.** High-entropy individuals performed significantly faster (a) accelerations, decelerations, and (b) rotations (black datapoints +/- 95% CI). However, these same performance metrics have little to no direct effect on moment-to-moment changes in entropy within individuals (red shaded areas show within-individual estimates). Note that for the translational maneuvers in (a), the within-individual effect is significant, but has a much smaller magnitude (weaker slope) than the corresponding among-individual estimates. A similar conclusion is drawn for the rotations (b), except that in this case the within-individual effect has a confidence interval that overlaps 0. The estimates in this figure are derived from unstandardized values to allow among- and within-individual slopes to be compared.

**Table S1. Number of maneuvers per 2-s window of forward flight.** Descriptive statistics are provided from the comparative study (n = 732,855 windows from 207 individuals).

| | **Mean (± SD)** | **Quantile** | | | | |
|---|---|---|---|---|---|---|
| | | **5th percentile** | **25th percentile** | **Median** | **75th percentile** | **95th percentile** |
| Number of translations | 2.45 (1.48) | 0 | 1 | 2 | 4 | 5 |
| Number of rotations | 0.70 (1.08) | 0 | 0 | 0 | 1 | 3 |
| Number of arcing turns | 0.23 (0.48) | 0 | 0 | 0 | 0 | 1 |
| Number of pitch-roll turns | 0.36 (0.58) | 0 | 0 | 0 | 0 | 1 |
| Number of distinct types of translations and rotations | 2.12 (0.93) | 1 | 2 | 2 | 3 | 4 |

**Table S2. Analysis of the repeatability of entropy.** Results are shown for a mixed-effects model analyzing trial-average entropy as a function of site and competitor presence, with a random effect of bird ID (n = 52 trials of 20 individuals).

| **Predictor** | **Estimate** | **95% confidence interval** | **t** | **p-value** |
|---|---|---|---|---|
| Site | | | | |
| California 1 | 0.80 | 0.74, 0.86 | -- | -- |
| California 2 | 0.85 | 0.81, 0.89 | -- | -- |
| Vancouver | 0.90 | 0.85, 0.94 | -- | -- |
| Competitor present vs. absent | 0.002 | –0.03, 0.03 | 0.13 | 0.90 |
| **Variance components** | **Variance** | | | |
| Bird ID | 0.002825 | | | |
| Residual | 0.002683 | | | |

**Table S3. Relationship between a bird's average maneuvering performance and its unpredictability.** Each model analyzed a bird's trial-average entropy as a function of trial-average performance on a specific maneuver, using data from the comparative study (n = 207 individuals from 25 species). The posterior mean is the slope estimated from the Bayesian regression model, accounting for field site as an additional fixed effect and with the phylogeny in the random effects structure. Entropy and performance values were standardized so that the effect sizes are comparable throughout the table. Note that the metrics $DecHor_{\mathrm{max}}$, $PitchD_{\mathrm{vel,avg}}$, $Arc_{\mathrm{rad}}$, and $PRT_{\mathrm{time}}$ were multiplied by –1, such that a positive slope indicates that entropy increases with more challenging performance.

| **Model number** | **Performance metric** | **Posterior mean** | **95% credible interval** | | **Effective sample size** | **pMCMC** |
|---|---|---|---|---|---|---|
| | | | **Lower** | **Upper** | | |
| 1 | $Vel_{\mathrm{max}}$ | –0.22 | –0.34 | –0.10 | 1,127 | < 0.001 |
| 2 | $AccHor_{\mathrm{max}}$ | 0.36 | 0.23 | 0.47 | 671 | < 0.001 |
| 3 | $DecHor_{\mathrm{max}}$ | 0.44 | 0.31 | 0.57 | 882 | < 0.001 |
| 4 | $PitchU_{\mathrm{vel,avg}}$ | 0.42 | 0.27 | 0.58 | 1,000 | < 0.001 |
| 5 | $PitchD_{\mathrm{vel,avg}}$ | 0.48 | 0.34 | 0.61 | 1,000 | < 0.001 |
| 6 | $Yaw_{\mathrm{vel,avg}}$ | 0.43 | 0.28 | 0.58 | 1,000 | < 0.001 |
| 7 | $Arc_{\mathrm{rad}}$ | –0.08 | –0.21 | 0.05 | 1,000 | 0.24 |
| 8 | $Arc_{\mathrm{vel,avg}}$ | 0.08 | –0.04 | 0.21 | 1,000 | 0.24 |
| 9 | $Arc_{\mathrm{cent,max}}$ | –0.04 | –0.16 | 0.10 | 1,133 | 0.55 |
| 10 | $PRT_{\mathrm{deg}}$ | 0.06 | –0.07 | 0.18 | 843 | 0.35 |
| 11 | $PRT_{\mathrm{time}}$ | –0.15 | –0.28 | –0.03 | 1,000 | 0.03 |
| 12 | $\%PRT$ | 0.39 | 0.26 | 0.52 | 1,000 | < 0.001 |

**Table S4. Relationship between biomechanical traits and unpredictability in the comparative study.** This model analyzed a bird's trial-average entropy as a function of muscle capacity and morphology, controlling for field site. Muscle capacity was measured at the species level, whereas body mass, wing loading, and aspect ratio were measured from the same individuals that were flown in the flight trials. The posterior mean is the slope estimated from the Bayesian phylogenetic regression model on unstandardized data (n = 207 individuals from 25 species).

| Trait | Posterior mean | 95% credible interval | | Effective sample size | pMCMC |
|---|---|---|---|---|---|
| | | Lower | Upper | | |
| Burst muscle capacity | 0.01 | –0.01 | 0.03 | 1,183 | 0.22 |
| Body mass (g) | –0.004 | –0.03 | 0.02 | 778 | 0.80 |
| Wing loading (g/cm$^2$) | –0.52 | –0.89 | –0.08 | 1,000 | 0.02 |
| Aspect ratio | 0.02 | –0.02 | 0.05 | 1,000 | 0.35 |

**Table S5. The effect of simulated changes in speed on unpredictability.** This analysis compared trial-average entropy for 20 individual hummingbirds at their original speed vs. two simulated speed conditions, where behavior was simulated to occur in either 2x faster or 2x slower. Results are shown for a mixed-effects model with bird ID as a random effect and speed condition as the predictor (n = 52 trials of 20 individuals). The estimate provides the average change in entropy, relative to the original trial.

| Speed condition | Estimate | 95% confidence interval | | t | p-value |
|---|---|---|---|---|---|
| | | Lower | Upper | | |
| 2x faster | 0.51 | 0.48 | 0.54 | 37.78 | < 0.0001 |
| 2x slower | –0.37 | –0.40 | –0.35 | –28.07 | < 0.0001 |

**Table S6. Within-individual analysis of unpredictability.** This analysis tested the influence of specific maneuvers on moment-to-moment changes in entropy, using data from the comparative study. Results are reported for a mixed-effects model analyzing 251,545 windows of flight (205 individuals total) that accounted for temporal autocorrelation; this sample represents windows that had at least one translation and one rotation maneuver. All predictors variables and the response variable were standardized so that the effect sizes are comparable throughout the table. As an additional measure of effect size, the table also provides the average Pearson's correlation statistics for the within-individual correlations (averaged across n = 88 individuals that had > 1,000 windows meeting the criteria above).

| **Predictor (standardized)** | **Estimate** | **95% confidence interval** | | **t** | **p-value** | **Within-individual correlation** | |
|---|---|---|---|---|---|---|---|
| | | **Lower** | **Upper** | | | **R** | **$R^2$** |
| Travel velocity | –0.41 | –0.42 | –0.40 | –86.38 | < 0.0001 | –0.27 | 8.9 % |
| Number of translations | –0.01 | –0.02 | –0.01 | –7.12 | < 0.0001 | –0.03 | 1.1 % |
| Number of rotations | 0.01 | 0.006 | 0.014 | 5.00 | < 0.0001 | 0.09 | 1.4 % |
| Number of arcing turns | 0.015 | 0.011 | 0.018 | 7.40 | < 0.0001 | –0.004 | 0.4 % |
| Number of pitch roll turns | 0.018 | 0.013 | 0.022 | 8.39 | < 0.0001 | –0.05 | 1.0 % |
| Translational performance | 0.085 | 0.08 | 0.09 | 29.24 | < 0.0001 | 0.08 | 1.7 % |
| Rotational performance | –0.003 | –0.008 | 0.002 | –1.10 | 0.27 | –0.01 | 0.5 % |
